# Cytopenias and Functional Defects in a Novel Murine Model of VPS45 Severe Congenital Neutropenia

**DOI:** 10.64898/2026.06.15.732420

**Authors:** Josias Soares de Brito, Zhiqing Zhu, Kristyn Norris, Natasha Buwa, Melonnie Furgason, Peter Opara-Nadi, Bruce Woda, Christoph Klein, Mary Munson, Peter E. Newburger

## Abstract

Mutations in the *VPS45* gene are associated with a rare form of severe congenital neutropenia (SCN5), a life-threatening inherited error of immunity. We developed and characterized a novel mouse model of SCN5 by CRISPR/Cas9-mediated knock-in of pathogenic *VPS45* E238K and T224N mutations. Both *Vps45* mutations led to decreased protein expression in bone marrow cells. *In vivo* phenotyping demonstrated a non-Mendelian genetic distribution with reduced numbers of knock-in homozygotes *Vps45*^E238K^. *Vps45*^E238K^ knock-in homozygous mice showed reduced body weight, reduced body condition with age, and increased mortality. As in human SCN5, *Vps45*^E238K^ knock-in homozygotes demonstrated neutropenia and lymphopenia. Functionally, *Vps45*^E238K^ knock-in homozygote neutrophils exhibited increased lipopolysaccharide-induced apoptosis and decreased peroxide production, phagocytic capacity and *in vivo* cell migration, phenocopying the functional defects reported in patients. *Vps45*^T224N^ knock-in homozygous mice showed a milder phenotype or no abnormalities. In conclusion, this mouse model phenocopies, in part, human SCN5. It provides a novel platform for future studies of the pathophysiology of defects in neutrophil number and function in human SCN5, potential therapies for the disease, and the biochemistry and cell biology of VPS45.

**Summary statement:** We report a mouse model of severe congenital neutropenia due to VPS45 missense mutations. It represents the first animal model of human neutropenia due to a defect in intracellular trafficking.

## Introduction

The endosomal-lysosomal vacuolar protein sorting regulator VPS45 is a conserved member of the Sec1/Munc18 family of proteins that regulate transport and assembly of SNARE complexes and SNARE-mediated fusion in all eukaryotes.(Zhang & Hughson 2021) In humans, mutations in VPS45 cause severe congenital neutropenia (SCN) type 5 (SCN5; OMIM # 615285). All reported pathogenic mutations are located in a putative hinge region of the protein, near a dominant negative mutation of yeast W244 (human W230) known to cause abnormal SNARE binding and accumulation of intracellular vesicles.(Carpp et al. 2006; Carr & Rizo 2010; Shah et al. 2017) Each of the human mutations likely alters the structure of this region of VPS45 leading to dysregulated SNARE complexes and fusion,(Carpp et al. 2006) which can decrease cell survival and function.

Two homozygous missense mutations, T224N and E238K, in conserved residues of VPS45 were first identified in several consanguineous families.(Stepensky et al. 2013; Vilboux et al. 2013) These two and additional novel VPS45 mutations, including P468L and R386H, have since been identified in unrelated kindreds in Israel and the U.S.(Shah et al. 2017; Shadur et al. 2019) Patients with these mutations suffer from recurrent life-threatening infections and die at age 3 months to 3 years if not treated by hematopoietic stem cell transplantation.(Shadur et al. 2019) All affected patients display granulocyte colony-stimulating factor (G-CSF)-unresponsive SCN, neutrophil and platelet dysfunction, and myelofibrosis. Some also have CD8 lymphopenia and defective T lymphocyte activation responses (Kostel Bal et al. 2024) or *Pneumocystis jirovecii* pneumonia suggestive of deficient T cell function (Shah et al. 2017). Several patients, from unrelated kindreds with E238K and P468L mutations, also have neurological or visual defects.(Shah et al. 2017; Karaatmaca et al. 2020; Vilboux et al. 2013; Stepensky et al. 2013; Meerschaut et al. 2015) (Kostel Bal et al. 2024).No other congenital anomalies have been reported.

Epstein-Barr virus-transformed lymphoblastic cell lines and fibroblasts from patients with VPS45 mutations demonstrated lower VPS45 levels, abnormally diffuse localization of VPS45, lower levels of VPS45 binding partners Syntaxin-16 and Rabenosyn-5, decreased lysosomes, decreased platelet α-granules, increased and enlarged mitochondria, slow migration, and accelerated apoptosis.(Stepensky et al. 2013; Vilboux et al. 2013) Frey *et al*. have demonstrated embryonic lethality in *Vps45* knockout (KO; *Vps45*^KO^) mice, with *in vitro* studies showing that loss of Vps45 leads to defective trapping of cargos in early endosomes and impaired delivery to lysosomes.(Frey et al. 2021) However, the functional and physiological consequences of pathogenic missense mutations have not previously been studied in any animal model.

Here, we present a mouse model of SCN caused by *Vps45* T224N and E238K missense mutations that recapitulates in part the human disease (Stepensky et al. 2013; Vilboux et al. 2013). It should provide a powerful platform for understanding the primary defects of VPS45 mutation independent of the secondary effects of infections and inflammation in human infants with SCN5. This knock-in mutant mouse model, the first for any human neutropenia associated with an intracellular trafficking defect, provides new insights into the consequences of these mutations to the impairment of VPS45 function. It presents a novel platform for investigating the molecular pathobiology of pathogenic mutations and potential therapeutics for SCN5.

## Results

### Decreased Vps45 protein levels

*Vps45*^T224N^ and *Vps45*^E238K^ homozygous total bone marrow (BM) nucleated cells showed decreased intracellular protein levels, detected by immunoblotting (**Figure 1**), as in the human disease.(Vilboux et al. 2013; Stepensky et al. 2013) Vps45 protein was also decreased in heterozygous *Vps45*^KO^ BM, as expected.

**Figure 1.**
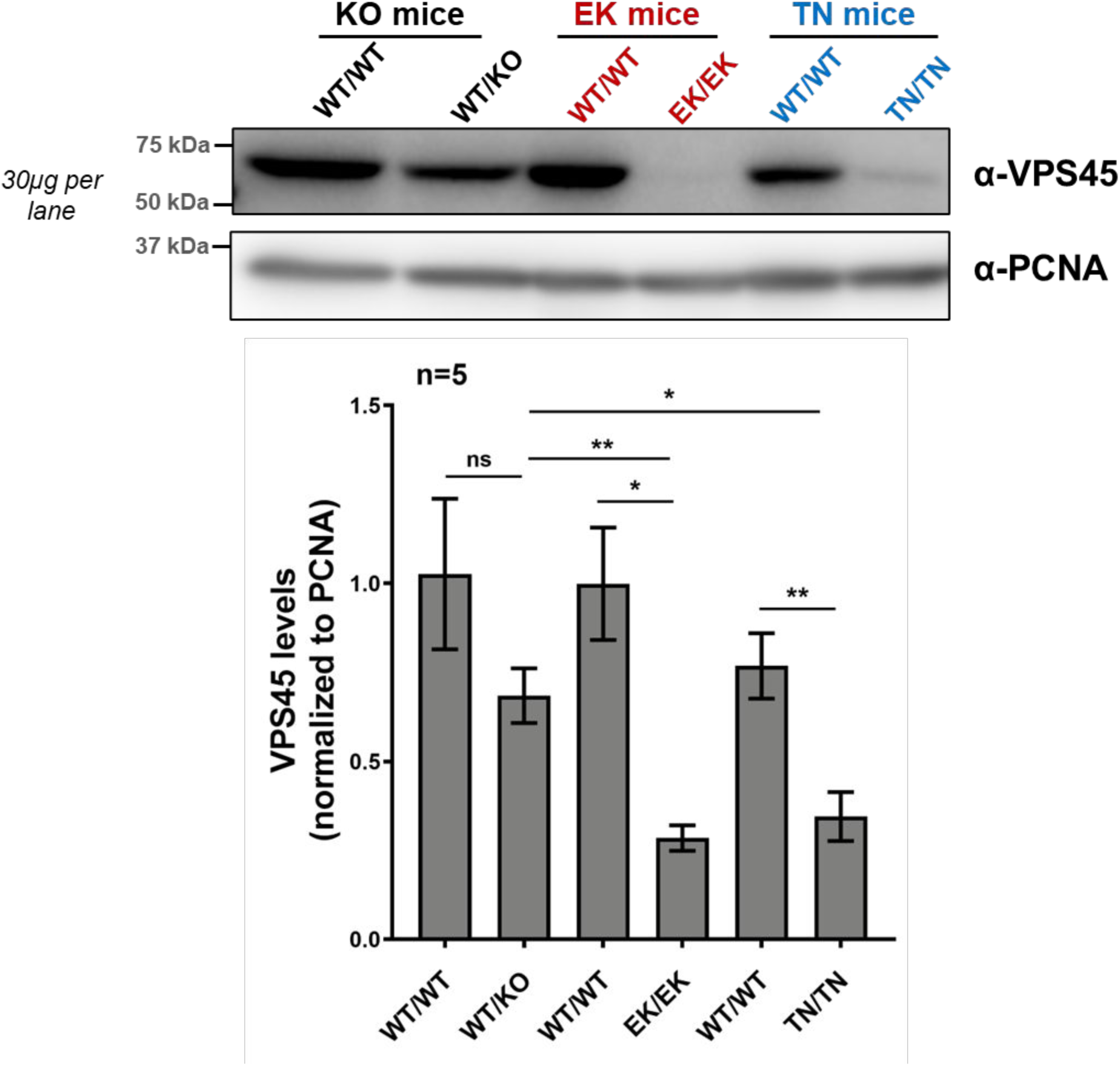
Immunoblot analyses of VPS45 expression in *Vps45* mice: *Vps45*^KO^-WT/KO, *Vps45*^E238K^-KI/KI and *Vps45*^T224N^-KI/KI mutations present reduced levels of Vps45 protein expression in BM cells. BM tissue from each genotype were lysed and Vps45 levels detected using α-VPS45 antibody. Band intensity corresponding to Vps45 was quantified using densitometry analysis and normalized to the respective PCNA control. Data are expressed as mean ± standard error from five independent experiments. Statistical analyses used paired two-tailed Student’s t-test, with *p* <0.05 considered significant.

### Vps45^E238K^ mice demonstrated reduced fitness

*Vps45*^E238K^ homozygous knock-in mutant mice of both sexes displayed significantly decreased mean body weights (18% to 34 % at 6- to 52-weeks) and visibly reduced body size compared to wild-type littermates (**Figure 2**); *Vps45*^T224N^ homozygote and *Vps45*^KO^ heterozygote mice presented no significant differences in body weight from wild-type littermates (**Figure 2**). However, *Vps45*^E238K^ homozygous mutant mice showed no significant organ weight differences in liver, spleen, heart, lungs, pancreas, or thymus when corrected for body weight (data not shown). We did not observe the nephromegaly characteristic of human VPS45 disease (Stepensky et al. 2013; Vilboux et al. 2013; Shah et al. 2017).

**Figure 2.**
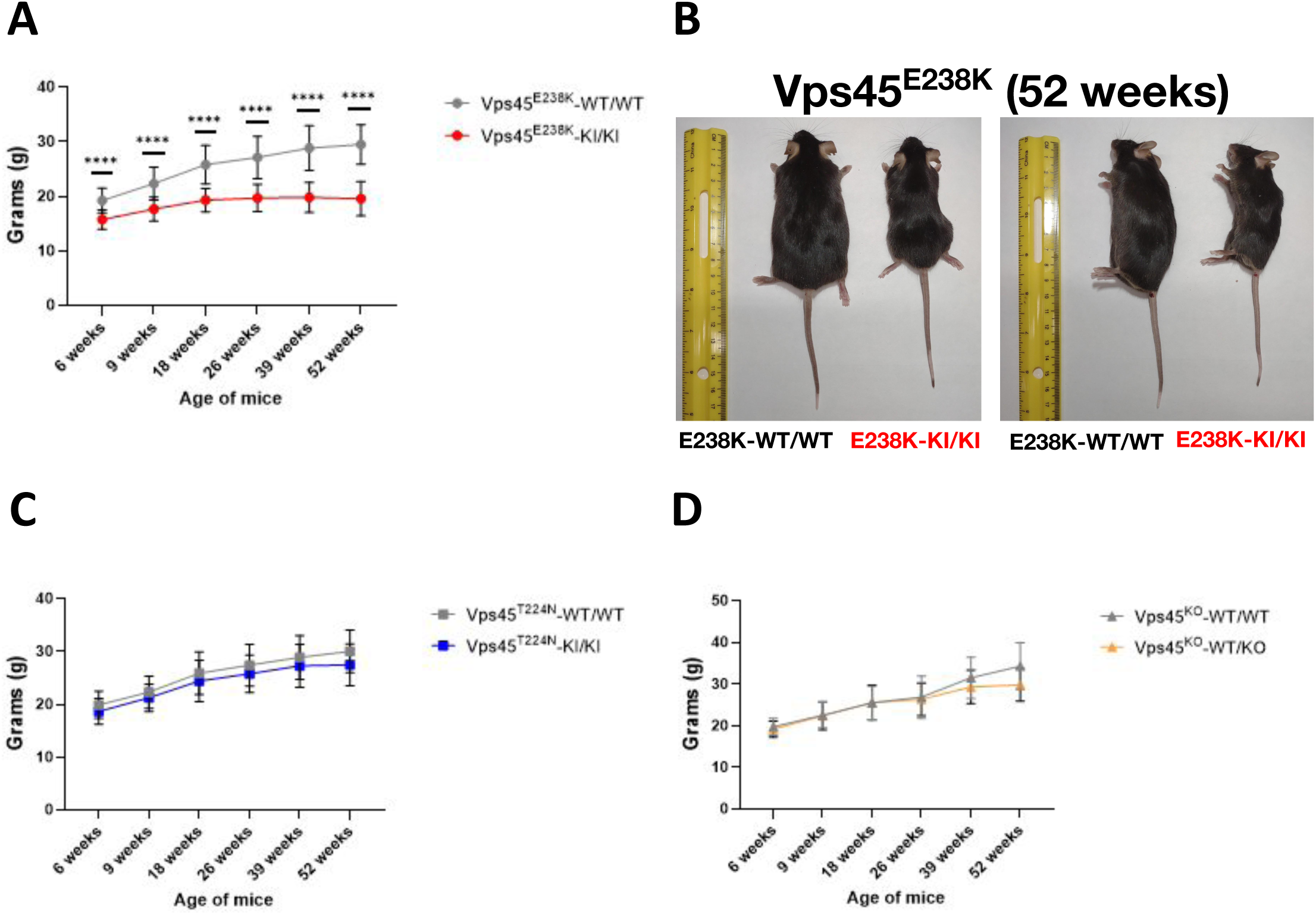
Body weight and size of *Vps45* mice: *Vps45* mouse body weight and size were evaluated from ages 6 to 52 weeks. **(A)** *Vps45*^E238K^-KI/KI showed reduced weight and size from 6 to 52 weeks of age; **(B)** Representative images of *Vps45*^E238K^-KI/KI with WT littermate at 52 weeks of age; **(C)** *Vps45*^T224N^-KI/KI displayed no significant change, and **(D)** no significant difference was observed in *Vps45*^KO^-WT/KO mice. Line graphs present mean ± standard deviation; unpaired, two-tailed Student’s t-tests were performed between the WT and KI mice of each genotype at each specific age, with *p*<0.05 considered significant. **** *p*<0.0001.

To investigate pre- and peri-natal fitness of mutant mice, we evaluated the genetic distribution of offspring from *Vps45*^T224N^ and *Vps45*^E238K^ heterozygote/heterozygote matings by DNA genotyping. *Vps45*^E238K^ offspring differed significantly (p-value < 0.0001, **\*\***) from a normal Mendelian distribution, with a decreased proportion of live offspring with the homozygous mutant genotype (**Figure 3**), with similar findings in both males and females. *Vps45*^T224N^ heterozygote/heterozygote and *Vps45*^KO^ heterozygote/wild-type matings both produced normal Mendelian distributions of offspring. Both *Vps45*^T224N^ and *Vps45*^E238K^ homozygous males were fertile. *Vps45*^E238K^ homozygous females could become pregnant but did not have observed post natal progeny; whereas, *Vps45*^T224N^ homozygous females produced offspring in expected numbers.

**Figure 3.**
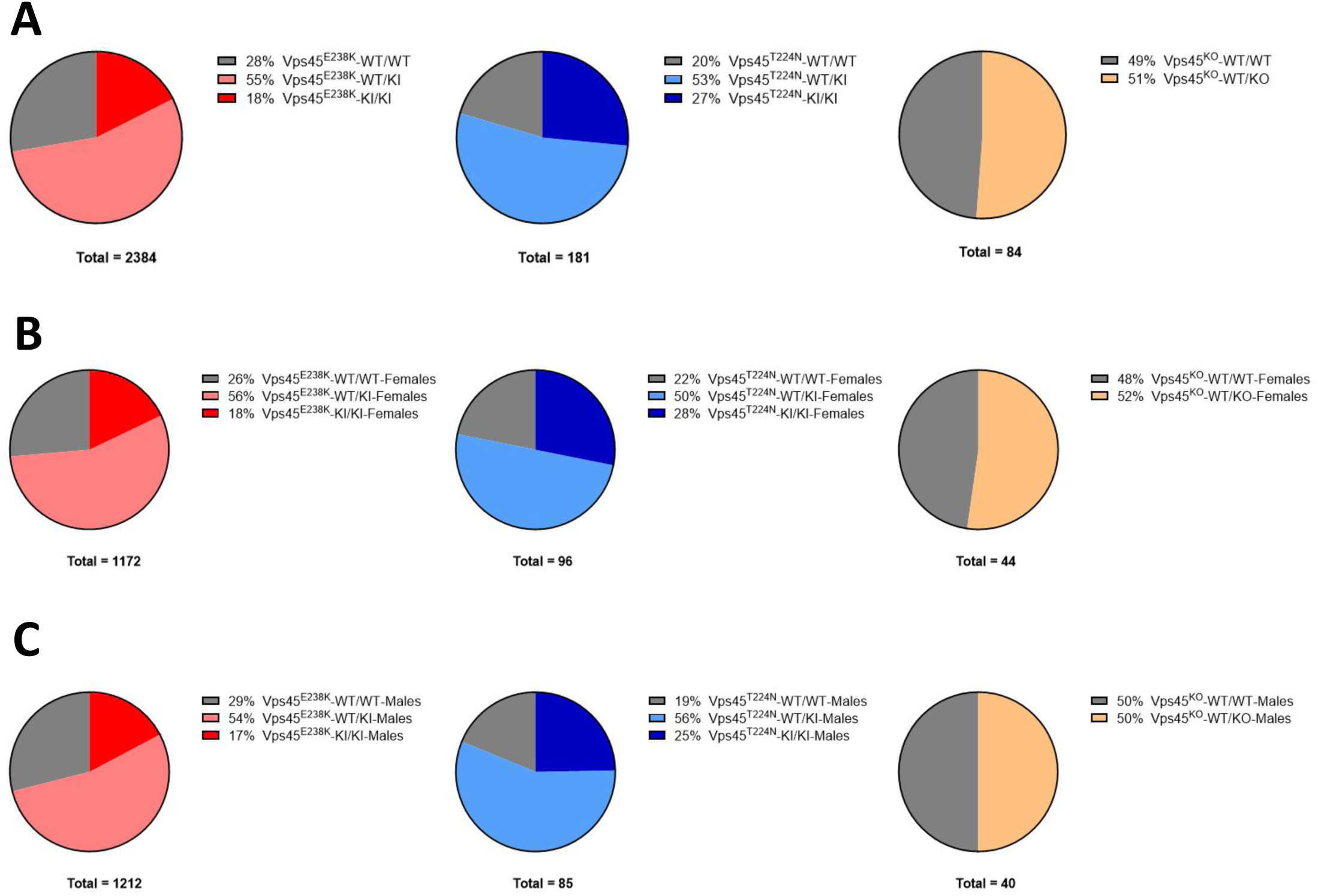
Mendelian genetic distribution of *Vps45* mice: Offspring from heterozygote crossings were genotyped at 3 weeks after birth and had their sex and genetic frequency evaluated for *Vps45*^KO^ (15 litters and 84 observed mice), *Vps45*^T224N^ (32 litters and 211 observed mice) and *Vps45*^E238K^ (380 litters and 2438 observed mice). Observed sex frequencies in *Vps45*^KO^, *Vps45*^T224N^ and *Vps45*^E238K^ follow the expected distributions. The observed genotype frequencies in *Vps45*^KO^ heterozygote/wild-type matings and *Vps45*^T224N^ heterozygote matings follow the expected distributions, while *Vps45*^E238K^ heterozygote matings do not follow a normal Mendelian distribution, either for all offspring **(A)** (p-value < 0.0001, ****) or for females **(B)** and males **(C)** separately analyzed. Chi-square test was performed, with *p*<0.05 considered significant.

To investigate fitness of the knock-in mutant mice with respect to aging, we assessed body condition and mortality over time. Measurements of body condition of *Vps45*^E238K^ homozygous mice up to age 1 year revealed decreased body condition scores compared to wild-type littermates (**Supplementary Figure 1**) and increased mortality in mice above age 26-weeks, with the effect seen largely in females (**Figure 4**).

**Figure 4.**
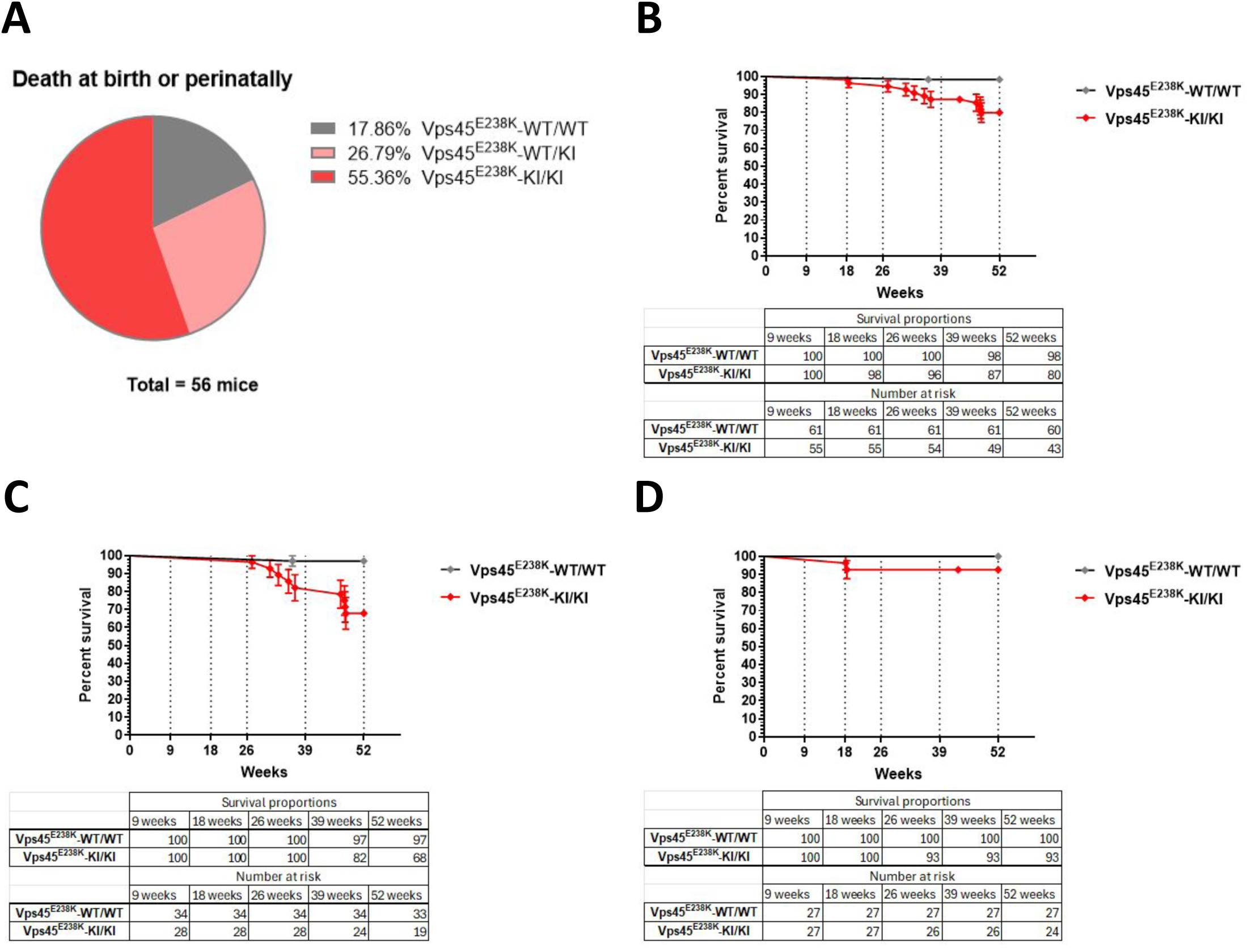
Survival of *Vps45* mice: *Vps45*^T224N^ and *Vps45*^E238K^ mice from heterozygote crossings showed increased mortality for *Vps45*^E238K^ homozygous mice, whereas *Vps45*^T224N^ homozygotes demonstrated a milder increase in mortality. **(A)** Proportions of *Vps45*^E238K^-WT/WT (10 mice), *Vps45*^E238K^-WT/KI (15 mice) and *Vps45*^E238K^-KI/KI (31 mice) that were observed dead at birth or perinatally (up to 6 weeks post-birth). **(B-D)** Kaplan-Meier survival curves as well as survival proportions and numbers at risk are represented for *Vps45*^E238K^-WT/WT and *Vps45*^E238K^-KI/KI **(B)**females and males, **(C)** females and **(D)** males. Kaplan-Meier estimates were analyzed by Mantel-Cox log-rank and Gehan-Breslow-Wilcoxon tests, with *p*<0.05 considered significant. Error bars represent standard errors of means. * *p*<0.05, ** *p*<0.01, *** *p*<0.001, **** *p*<0.0001.

### Vps45^E238K^ mutation causes decreased neutrophil, monocyte, lymphocyte, and platelet numbers

Peripheral blood cell counts in *Vps45*^E238K^ homozygous mutant mice at age 9-weeks showed significantly reduced white blood cell counts with significant reductions in absolute numbers of lymphocytes, monocytes and neutrophils. Hemoglobin levels were similar between mutant and wild-type mice. *Vps45*^E238K^ homozygote mice also showed significantly decreased platelet counts (**Figure 5 and Supplementary Figure 2**) with increased platelet distribution width and coefficient of variation of the platelet distribution width (**Supplementary Figure 2**).

**Figure 5.**
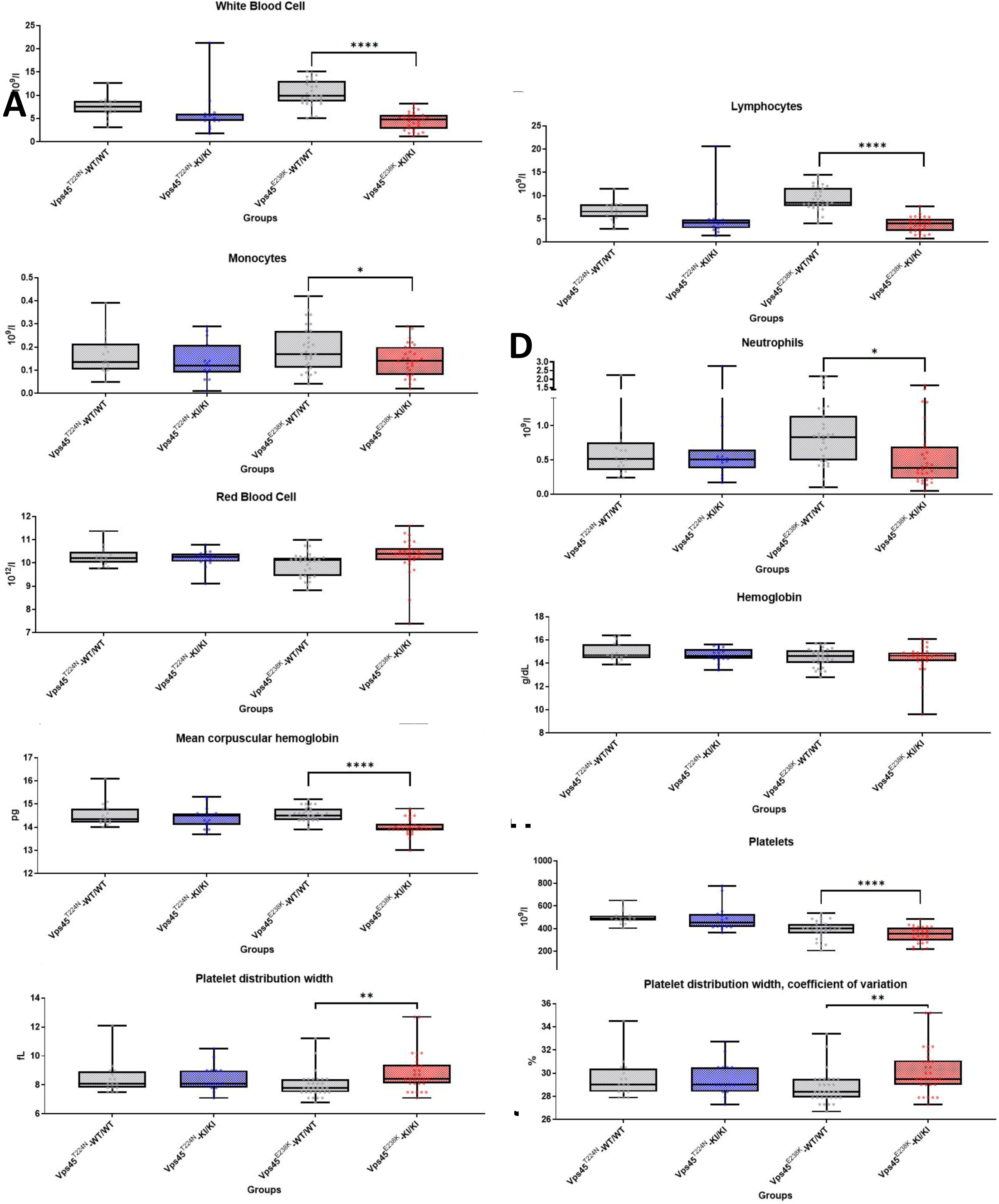
Complete blood counts of *Vps45* mice at 9 weeks of age: Peripheral blood CBC of *Vps45* mice at 9 weeks of age demonstrate decreases in **(A)** white blood cells, **(B)** lymphocytes, **(B)**monocytes, **(D)** neutrophils, **(E)** red blood cells, **(F)** hemoglobin, **(G)** mean corpuscular hemoglobin, **(H)** platelets, **(I)** platelet distribution width and **(J)** platelet distribution width coefficient of variation for *Vps45*^E238K^-KI/KI relative to *Vps45*^E238K^-WT/WT with no significant differences observed in *Vps45*^T224N^ and *Vps45*^KO^ lines; hemoglobin levels did not differ in 9 week old mice. Box-and-whisker plots show minimum, interquartile range, median and maximum. Unpaired, two-tailed Student’s t-tests were performed between the WT and mutated mice of each genotype at 9 weeks of age, with *p*<0.05 considered significant. See **Supplementary Figure 3** for complete blood counts of *Vps45* mice from 6 to 52 weeks.

Homozygous *Vps45*^E238K^ mice also showed increases of both neutrophil:lymphocyte and platelet:lymphocyte ratios,(Hickman 2017; Neumann 2021) most likely resulting from the profound lymphopenia (data not shown). In contrast, all cell counts of *Vps45*^KO^ heterozygous mice were similar to those of wild-type littermates (**Figure 5 and Supplementary Figure 2**). Flow cytometry analysis of peripheral blood leukocytes and splenocytes of *Vps45*^E238K^ homozygous mice (**Figure 6**) confirmed reductions in peripheral blood and splenic neutrophils. Lymphocyte subtyping in the spleen showed profound decreases in CD4^+^, CD8^+^, and double-negative (CD4^-^, CD8^-^) T-lymphocytes but no significant differences in B-lymphocytes or NK cells (**Figure 6**).

**Figure 6.**
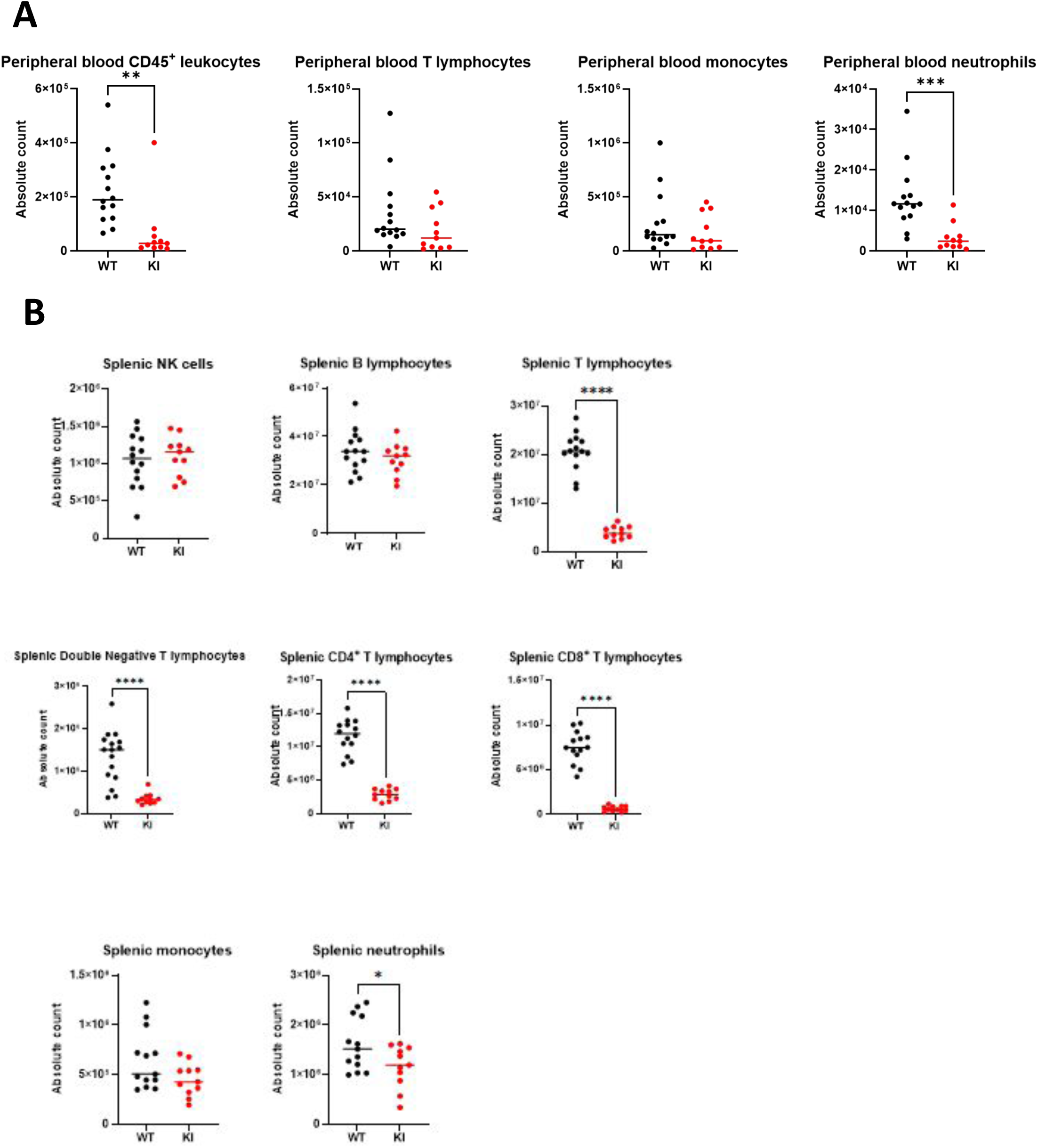
Immunophenotyping of blood and spleen leukocytes of *Vps45* mice: Flow cytometric analysis of peripheral blood and spleen leukocytes from 9 week-old *Vps45*^E238K^-WT/WT and *Vps45*^E238K^-KI/KI mice (10 mice for peripheral blood; 11 mice for spleen) confirmed CBCs results by demonstrating **(A)** decreased peripheral blood neutrophils, **(B)** decreased splenic T (CD3^+^/B220^-^) lymphocytes, T (CD4^-^/CD8^-^) double-negative lymphocytes, T CD4^+^ lymphocytes and T CD8^+^ lymphocytes, but no significant changes in splenic NK (CD122^+^/NK1.1^+^) cells and B (CD3^-^/B220^+^) lymphocytes. Scatter plots present medians with all values represented and p-values of Welsh’s parametric unpaired, two-tailed t-test between *Vps45*^E238K^-WT/WT and *Vps45*^E238K^-KI/KI; p-value <0.05 was considered significant. * *p*<0.05, ** *p*<0.01, *** *p*<0.001, **** *p*<0.0001.

### Vps45 mutant mice showed no differences in histology or cell ultrastructure

Given the observation of neutropenia and lymphopenia and the reported abnormal BM morphology (myelofibrosis) in patients, we examined mouse tissues for possible abnormalities, However, no significant histological changes were observed in thymus or BM of *Vps45*^T224N^ and *Vps45*^E238K^ homozygous mice (**Supplementary Figure 3**), nor in liver, spleen, heart, or kidneys (data not shown). Transmission electron microscopy of lymphocytes, neutrophils and platelets (**Supplementary Figure 4**) revealed no ultrastructural differences between cells from *Vps45*^E238K^ homozygous mutant and wild-type mice.

Myelofibrosis, a key feature of the human disease,(Vilboux et al. 2013; Stepensky et al. 2013) was not detectable histologically in 9 or 52 week-old *Vps45*^E238K^ homozygote mutant mice (**Supplementary Figure 3**). RT-qPCR for *Col3A1* as a molecular marker for myelofibrosis did not show any significant difference in expression between *Vps45*^E238K^ homozygote mutant and wild-type BM at age 52-weeks (**Supplementary Figure 5**).

### Apoptosis in Vps45 mutant mouse neutrophils

Increased apoptosis of mature neutrophils or myeloid precursors is a common mechanism of many forms of human congenital neutropenia, including VPS45 neutropenia.(Glaubach et al. 2014; Vilboux et al. 2013; Stepensky et al. 2013) We assessed mature peripheral blood neutrophils for apoptosis by flow cytometric analysis of annexin-V binding and PI content for the detection of early (annexin-V^+^/PI^-^) and late (annexin-V^+^/PI^+^) apoptotic events. No significant differences were observed under spontaneous conditions (data not shown). Upon exposure to bacterial LPS, neutrophils from *Vps45*^T224N^ and *Vps45*^E238K^ homozygotes did not show significant differences in numbers of late or early apoptotic cells compared to wild-type littermates (**Figure 7A**). However, neutrophils from both knock-in mutant genotypes when gated for live cells (PI negative events) showed significantly increased binding (MFI) of annexin-V (**Figure 7B**).

**Figure 7.**
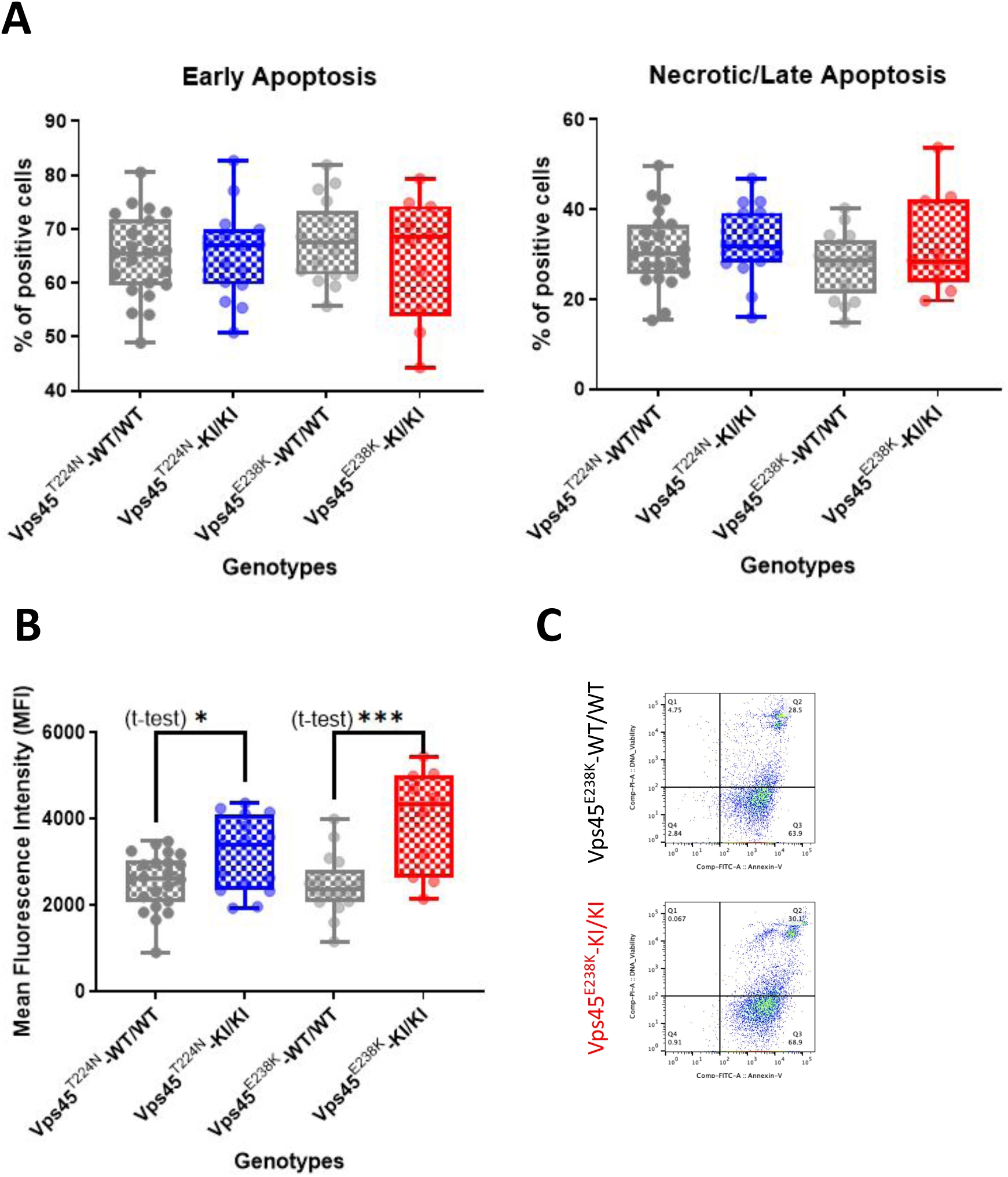
Neutrophil apoptosis in *Vps45* mice: Bacterial lipopolysaccharide increased binding of annexin-V in *Vps45*^T224N^-KI/KI and *Vps45*^E238K^-KI/KI mice. Blood leukocytes of 9 week-old *Vps45*^T224N^ and *Vps45*^E238K^ homozygote mice were incubated with lipopolysaccharide for 2 hours, stained for neutrophil cell surface markers (CD11b and Ly6G) and stained for the detection of annexin-V and propidium iodide (PI) for measurements of apoptosis by flow cytometry. **(A)** Flow cytometric analysis of CD11b^+^Ly6G^+^ blood leukocytes for early (annexin-V^+^/PI^-^) and late (annexin-V^+^/PI^+^) apoptosis demonstrates no significant difference between *Vps45*^T224N^-KI/KI (15 mice) and *Vps45*^E238K^-KI/KI (10 mice) compared to WT littermates (17 *Vps45*^E238K^-WT/WT mice and 22 *Vps45*^T224N^-WT/WT mice). **(B)** MFI values of annexin-V in live (PI^-^)/CD11b^+^Ly6G^+^ blood leukocytes demonstrate increased expression of phosphatidylserine, a marker of apoptosis, on both genotypes *Vps45*^T224N^ (*p* = 0.02) and *Vps45*^E238K^ (*p* = 0.0004) by unpaired, two-tailed Student’s t-test. **(C)** Representative *Vps45*^E238K^-WT/WT and *Vps45*^E238K^-KI/KI flow plots of annexin-V expression on gated live CD11b^+^Ly6G^+^ double positive cells. Box-and-whisker plots present minimum, interquartile range, median and maximum with p-values of t-test between groups based on values obtained from at least nine experiments; p-value <0.05 was considered significant. * *p*<0.05, *** *p*<0.001. See **Supplementary Figure 7** for gating strategy.

### Vps45^E238K^ neutrophils display impaired function

The normal function of neutrophils in host defense against infection includes migration to sites of microbial invasion, ingestion of microbes (phagocytosis), and killing of microbes by an array of “weapons,” the most essential of which is generation of hydrogen peroxide by the phagocyte NADPH oxidase complex (Siwicki & Kubes 2023). As neutrophils from VPS45 patients have been shown to have deficient peroxide production and motility,(Vilboux et al. 2013) we evaluated mature peripheral blood neutrophils in our *Vps45* knock-in mice for their capacity to produce peroxide (**Figure 8A**) by measurement of DHR fluorescence upon activation. PMA-stimulated peroxide production was significantly decreased in neutrophils from *Vps45*^E238K^ homozygote mice. Furthermore the phagocytic capacity of neutrophils from *Vps45*^E238K^ homozygote mice was decreased, both in the percentage of cells ingesting fluorescently-labeled *E. coli* (**Figure 8B**, left panel) and numbers of bacteria ingested and phagosome acidification (MFI in **Figure 8B**, right panel). No differences were observed when neutrophils were challenged with either *S. aureus* or zymosan particles.

**Figure 8.**
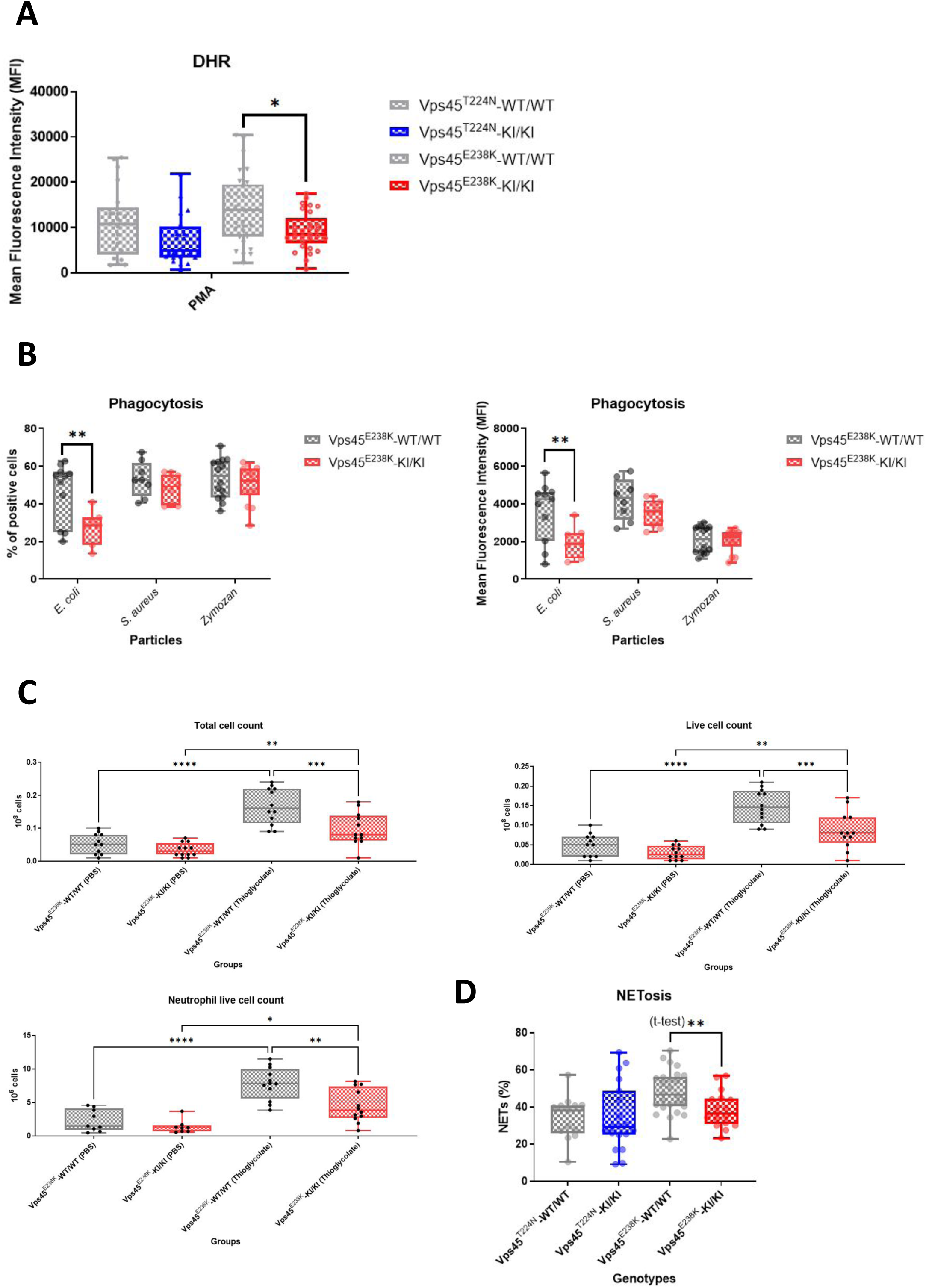
Neutrophil function in *Vps45* mice: *Vps45*^E238K^-KI/KI mice presented reduced oxidative burst activity, phagocytosis of *E. coli* particles, *in vivo* cell migration and NETosis. **(A) DHR fluorescence.** Flow cytometric detection of PMA-stimulated DHR fluorescence revealed statistically significant differences between DHR MFI signal from neutrophils of *Vps45*^E238K^-KI/KI and *Vps45*^E238K^-WT/WT littermates (*p* = 0.02; one-way ANOVA with Tukey’s multiple comparisons test). **(B) Phagocytosis.** Percentage (left panel) and MFI (right panel) of neutrophils incubated with fluorescently labeled *E. coli*, *S. aureus* and zymosan particles demonstrated significant difference in % positive cells (*p* = 0.002) and MFI of pooled cells (*p* = 0.003) by two way ANOVA with Šidák’s multiple comparisons test. **(C) *In vivo* migration**. Leukocytes harvested from mouse peritoneal cavities after phosphate-buffered saline (PBS) or thioglycollate injections were assessed for total cell (left panel), live cell (middle panel) and neutrophil (right panel) counts. All three showed significant increases in cell counts in thioglycolate compared to PBS injections. Significant differences were observed between *Vps45*^E238K^-KI/KI and *Vps45*^E238K^-WT/WT littermates in total cell count (*p* = 0.0009), live cell count (*p* = 0.001), and live neutrophil count (*p* = 0.002 by one way ANOVA with Tukey’s multiple comparisons test). **(D) NETosis.** Flow cytometric detection of Hoechst 33342 (DNA), myeloperoxidase (MPO) and citrullinated histone H3 in PMA-stimulated *Vps45*^T224N^ and *Vps45*^E238K^ mouse neutrophils demonstrated no significant difference between *Vps45*^T224N^-KI/KI and *Vps45*^T224N^-WT/WT and a significant difference between *Vps45*^E238K^-KI/KI and *Vps45*^E238K^-WT/WT (*p* = 0.006 by Student’s t-test). Box-and-whisker plots present minimum, interquartile range, median and maximum from nine or more experiments. In all experiments, a *p* <0.05 was considered significant. * *p*<0.05, ** *p*<0.01, *** *p*<0.001, **** *p*<0.0001. See **Supplementary Figure 7** for gating strategy.

Neutrophil *in vivo* cell migration, which is impaired in human VPS45 deficient neutrophils,(Vilboux et al. 2013) was assessed by measuring transmigration into the peritoneum induced by thioglycolate(**Figure 8C**). Peritoneal leukocyte numbers were similar for viable resident cells after PBS injection, but significantly decreased in thioglycolate-elicited peritoneal exudates in *Vps45^E238K^* homozygote mice (**Figure 8C**). Total cell counts with leukocyte differential counts showed lower numbers of neutrophils in the thioglycolate-elicited peritoneal exudates (**Supplementary Table 1**).

The capacity of neutrophils to undergo NETosis upon PMA stimulation, a function generally dependent upon both granule mobilization and peroxide production,(Kenny et al. 2017) was evaluated by flow cytometry for cells staining for DNA (Hoechst 3342), citrullinated histone H3 (citH3), and myeloperoxidase (MPO) (**Figure 8D**). NETosis was significantly decreased in neutrophils from *Vps45*^E238K^ homozygotes.

## Discussion

We report the first mouse model of VPS45-associated SCN, an inherited error of immunity, using human pathogenic mutations inserted into the mouse *Vps45* gene using CRISPR/Cas9 gene editing. Our findings indicate that homozygosity for the *Vps45*^T224N^ and *Vps45*^E238K^ mutations results in decreased Vps45 protein expression in BM cells (**Figure 1**), consistent with reports in SCN5 patients’ EBV-transformed lymphoblast cell lines and fibroblasts.(Vilboux et al. 2013; Stepensky et al. 2013) The resultant mouse phenotype featured an imbalance in the Mendelian genetic distribution in the *Vps45*^E238K^ mouse line, with fewer knock-in homozygotes than expected (**Figure 3**). Possible intrauterine impairment of embryonic development and/or early postnatal mortality might underlie the skewed ratios. Reduced capacity to thrive was also evident from smaller body size and weight in adult mice (**Figure 2**), with health impairment with aging evident by decreasing body condition scores (**Supplementary Figure 1**) and increased late mortality (**Figure 4**). The long-term effects of human VPS45 mutations remain unknown, as all reported patients have died or received hematopoietic stem cell transplants by age 3 years.(Shadur et al. 2019; Vilboux et al. 2013; Stepensky et al. 2013; Karaatmaca et al. 2020) The mouse model also provides insights into non-hematopoietic phenotype features that would not be corrected by transplant.

Analysis of peripheral blood cells (**Figure 5**, **Figure 6 and Supplementary Figure 2**) demonstrated decreased numbers of lymphocytes, neutrophils, and platelets in homozygous *Vps45*^E238K^ mice, suggesting a lymphoid and myeloid multi-lineage effect of this point mutation. The lower platelet counts, with increased variation in platelet distribution width and its coefficient of variation (**Figure 5 and Supplementary Figure 2**), suggested stress in that lineage as well. We did not observe a decrease in platelet granules (**Supplementary Figure 4**), which is inconsistent in human SCN5 and might be a secondary effect of inflammatory signalling.

VPS45-deficient patient neutrophils demonstrate accelerated spontaneous apoptosis, which could be a primary mechanism responsible for cytopenias or a secondary effect of chronic infection.(Vilboux et al. 2013; Stepensky et al. 2013) In our study we found no spontaneous apoptosis, but neutrophils exposed to bacterial lipopolysaccharide showed a significant increase in annexin V binding (**Figure 7**), although that could also reflect a change in membrane phospholipid distribution due to a defect in endosomal trafficking. This finding suggests that the mutant neutrophils are more susceptible to undergoing cell death upon contact with gram-negative bacteria, and likely other micro-organisms, as may occur in infected children with *VPS45* defects.

Functional defects of *Vps45*^E238K^ homozygote neutrophils that phenocopied the human disease (Vilboux et al. 2013; Stepensky et al. 2013) included impairment in peroxide production (**Figure 8**), phagocytosis of *E. coli* (**Figure 8**), and *in vivo* cell migration (**Figure 8**), measured as thioglycolate-elicited peritoneal exudate formation. As the functional defects were relative rather than absolute, we did not test resistance to infection in these mice. Complete, or nearly complete, loss of function is necessary for a phenotype of increased infections in disorders of neutrophil function such as chronic granulomatous disease (Staudacher & von Bernuth 2024) and leukocyte adhesion deficiency.(Alasmari et al. 2023)

Flow cytometric analysis of lymphocyte subtypes (**Figure 6**) demonstrated that the lymphopenia observed in *Vps45*^E238K^ homozygous mouse blood counts consisted of deficiencies in CD4^+^, CD8^+^ and double-negative T lymphocytes, indicating a fundamental defect in T lymphocyte development. Profound T lymphopenia has been reported in one VPS45^E238K^ patient,(Karaatmaca et al. 2020) and another had reduced absolute counts of CD4^+^ and a more pronounced reduction of CD8^+^ lymphocytes and defective activation response to phytohemagglutinin and anti-CD3 antibody concomitant to profound neutropenia.(Kostel Bal et al. 2024) An additional patient with a P246L mutation developed *Pneumocystis jirovecii* pneumonia, indicating a likely T lymphocyte deficiency.(Shah et al. 2017)

Transmission electron microscopic imaging of *Vps45*^E238K^ mouse neutrophil, lymphocyte, and platelet morphology (**Supplementary Figure 4**) demonstrated no significant loss of cytoplasmic granules. Human VPS45^T224N^ patients’ neutrophils and platelets have been reported to have reduced granule contents,(Stepensky et al. 2013) although it is not known whether that is a primary defect or a secondary effect of degranulation with chronic inflammation and infection; a patient with VPS45^P246L^ had normal platelet granulation.(Shah et al. 2017)

Myelofibrosis is a hallmark of the human disease(Stepensky et al. 2013; Vilboux et al. 2013; Shah et al. 2017), also observed in studies utilizing BM-like organoids as a hematopoietic model to recapitulate the human VPS45 deficiency (Frey et al. 2021). However, histologic features of myelofibrosis were not detectable in young or old *Vps45*-mutant mice (**Supplementary Figure 3**). In addition, no significant histological alterations were observed in organs that showed extramedullary hematopoiesis in the VPS45-deficient patients, likely secondary to myelofibrosis.(Stepensky et al. 2013; Vilboux et al. 2013; Shah et al. 2017) These observations suggest a species difference in release of cytokines (e.g. transforming growth factor beta) or stability of megakaryocyte intracellular granules.

Our findings of defects in neutrophil function, without alterations in cellular morphology, suggest that function of these cells may be decreased due to perturbation of Vps45-mediated regulation of SNARE-dependent endosomal membrane trafficking by the structural alteration of the Vps45 hinge region. Another plausible hypothesis is that decreased stability of mutant Vps45 protein impacts its abundance while still preserving some normal protein functions. However, we did not find any abnormalities in cell counts or functions in heterozygous *Vps45*^KO^ mice with diminished Vps45 protein expression.

There are still very few mouse models of neutropenia that can be utilized for the study of new therapies. A mouse model of human SCN using patient-derived mutations in the GFI1 transcription factor showed a block in neutrophil differentiation as in the human disease (Zarebski et al. 2008). Mice genetically deficient in glucose-6-phosphatase-β display neutropenia with defective oxidative burst, calcium flux and chemotaxis functions.(Cheung et al. 2007) The only mouse model of neutropenia due to a defect in intracellular trafficking, the beige mouse model of Chediak Higashi syndrome due to *Lyst* mutations (Gallin et al. 1974), has a phenotype more dependent on impaired NK and cytotoxic T lymphocyte function (Roder 1979). Importantly, a mouse model of the most common genetic form of SCN due to mutations in the *ELANE* gene does not have a neutropenic phenotype (Grenda et al. 2002).

Our mouse model of *Vps45* missense mutations shares many of the pathophysiological characteristics of the human VPS45 SCN with consequences in the numbers of peripheral blood neutrophils and lymphocytes as well as impairments in several immunological functions of neutrophils, with impacts on both perinatal and long-term survival of these mice. The model should serve as a platform for future development of therapies for the human disease. Our *Vps45* mutant mice could also be used for studies of the biochemical, and cell biological consequences of *Vps45* mutations on endosomal-lysosomal trafficking.

## Materials and Methods

### Mice

Both mouse lines, *Vps45*^T224N^ and *Vps45*^E238K^, were generated by the UMass Chan Medical School Transgenic Animal Modeling Core. Mice were housed and bred in a specific pathogen-free environment. The study was conducted with approval of the Institutional Animal Care and Use Committee.

### Vps45 knock-in mice

The *Vps45*^T224N^ knock-in was created by blastocyst injection of C57BL/6 embryonic stem cells with CRISPR/Cas9-mediated gene editing to introduce the *Vps45* T224N mutation from a single-stranded donor (ss)DNA template and subsequent breeding of mice with germline insertions. Direct oocyte injections of Cas9/guide-RNA plasmid and *Vps45* E238K ssDNA template led to direct generation of *Vps45*^E238K^ mutant founder mice. Littermate homozygous knock-in (KI/KI) mutant, heterozygous (WT/KI), and wild-type (WT/WT) mice derived from gene-edited *Vps45*^T224N^ and *Vps45*^E238K^ founders were used in subsequent experiments after back-crossing to the C57BL/6J parental line for more than 7 (*Vps45*^T224N^) or 10 (*Vps45*^E238K^) generations for the elimination of off-target sites and mosaicism. All experiments used *Vps45*^T224N^ edited mice from a single founder; most experiments with *Vps45*^E238K^ mice used progeny from a single founder and were verified with repeat experiments with progeny of a second, independent founder. The CRISPR/Cas9 strategy, genomic maps, and sequence data are in **Supplementary Figure 6**; gRNA and oligonucleotide donor ssDNA sequences are in **Supplementary Table 2**.

### Vps45 knockout heterozygous mice

*Vps45*^KO^ heterozygous mice were generated from embryonic stem cells using a targeting vector to insert a trapping cassette in the intron upstream of exon 4 of *Vps45 (Frey et al. 2021)* using the “knockout-first” strategy. Frozen sperm of *Vps45*^KO^ heterozygous mice were used to re-generate the mouse line for our studies, and experiments were performed with heterozygous KO (WT/KO) and wild-type (WT/WT) littermate mice after 7 back-crosses with C57BL/6J mice.

### Mouse genotyping

Mouse genotyping was performed using PCR amplification of genomic DNA followed by restriction digestion of PCR products, with AseI for *Vps45*^T224N^ or HindIII for *Vps45*^E238K^ and without restriction digestion of PCR products for *Vps45*^KO^ (maps, methods, PCR primer, restriction digestion cut sites sequences in **Supplementary Methods, Supplementary Table 2** and **Supplementary Figure 6**. Sanger sequencing was performed to confirm the mutations (**Supplementary Figure 6.**

### Immunoblots

Total cell lysates of BM leukocytes were performed for the measurement of Vps45 protein expression. Lysates of *Vps45^KO^*HeLa cells (Frey et al. 2021) were used to confirm specificity of the anti-Vps45 antibody. Guinea pig anti-VPS45 polyclonal antibody was generated by Pocono Rabbit Farm & Laboratory (Canadensis, PA, USA; https://www.prfal.com/) using multiple injections of a purified recombinant human His_6_-tagged VPS45 protein (amino acids 1-556; 97% sequence homology and 99.5% sequence similarity between human and mouse orthologs), and affinity-purified using the recombinant VPS45(1-556)-His_6_ protein. Details are presented in **Supplementary Methods.** The anti-VPS45 primary antibody and horseradish peroxidase (HRP)-conjugated goat anti-guinea pig secondary antibody (Thermo Fisher Scientific, 61-4620) was used for Vps45 detection. Proliferating cell nuclear antigen (PCNA) served as a loading control, detected using a mouse monoclonal anti-PCNA antibody (Abcam, ab29) and HRP-conjugated goat anti-mouse secondary antibody (Jackson Immuno Research, 115-035-003). Immunoblots were prepared by standard methods (Taylor et al. 2013) with 30 µg of total protein from each cleared tissue lysate run on 10% SDS-PAGE gels and transferred to polyvinylidene difluoride membranes, which were probed by the indicated antibodies blocked with PBS/0.1% Tween/5% milk.

### Body weight and complete blood counts

Mice at ages 6– to 52-weeks were weighed, and peripheral blood samples collected from tail veins into EDTA blood collection tubes. Complete blood counts (CBCs) were determined by the VetScan HM5 (Abaxis) fully automated hematology analyzer and the haematological data from tail vein peripheral blood samples.

### Histology

Mice at ages 9- and 52-weeks had thymus, heart, lung, liver, spleen, pancreas, and kidney isolated, weighed, fixed in formalin and embedded in paraffin. Bones were similarly processed with an additional step for decalcification. Thin sections were stained with hematoxylin and eosin. For myelofibrosis detection, BM sections were stained with Gomorri’s trichrome and reticulin to assess the presence of BM fibrosis. All slides were interpreted by a pathologist (B.W.) highly experienced in mouse tissue histology. To confirm histochemistry results, molecular detection of myelofibrosis on BM and femoral bone cells was performed by real time quantitative PCR (RT-qPCR) detection of gene expression of *Col3A1* and actin alpha 2 (*Acta2)*, quantitated by 2^-ΔΔCt^ fold change; details are presented in **Supplementary Methods.**

### Electron microscopy

For evaluation of platelet ultrastructure, mouse peripheral blood was drawn by cardiac puncture and prepared as previously described (Koupenova et al. 2019) with minor modifications. Platelet pellets were stained and sectioned by standard methods and imaged on a Philips CM10 electron microscope by the UMass Chan Electron Microscopy Core.

Bone marrow neutrophils, isolated by negative magnetic enrichment using EasySep™ Mouse Neutrophil Enrichment Kit (STEMCELL Technologies, 19762) and CD3^+^CD8^+^ spleen cells, isolated by EasySep™ Mouse CD8^+^ T Cell Isolation Kit (STEMCELL Technologies, 19853), were fixed, stained, sectioned, and similarly imaged by standard methods by the UMass Chan Electron Microscopy Core.

### Flow cytometry

Flow cytometric immunophenotyping of leukocytes from peripheral blood and spleen tissue was performed to validate peripheral blood CBCs data and characterize peripheral blood and spleen CD45^+^ leukocytes, including splenic T lymphocyte subsets, B cells and NK cells (additional details in **Supplementary Methods**).

Flow cytometric analyses of apoptosis were performed using peripheral whole blood leukocytes subjected to two rounds of hypotonic red blood cell lysis in ice-cold water normalized to 1 x Dulbecco’s phosphate buffered saline (DPBS), centrifuged at 300 x g for 10 minutes and supernatant discarded. Cells were then subjected to Fc-block (BD Pharmingen™, 553142; 1:50) at 4°C for 5 minutes, stained with APC-R700 anti-CD11b (BD Horizon™, 564985; 3:100) and BV510 Ly6G antibodies (BD OptiBuild™, 740157; 3:100) at 4°C for 30 minutes for neutrophil identification and washed once in DPBS. Annexin V binding and PI uptake were assessed using the FITC Annexin-V Conjugate for Apoptosis Detection Kit (ThermoFisher, A13199) and PI (ThermoFisher, P3566) by following the manufacturer’s methods and taken for analysis by flow cytometry (BD Celesta). In experiments where bacterial LPS (300 ng/mL) was used to induce apoptosis, a two hour incubation was applied to the peripheral whole blood leukocytes cultivated in RPMI media without phenol red (ThermoFisher, 11835055) with 10% FBS and penicillin/streptomycin (Corning, 30-002-CI). CD11b^+^/ Ly6G^+^ gated mature neutrophils were evaluated for the frequency of annexin-V^+^/PI^-^(early apoptosis) and annexin-V^+^/PI^+^ (late apoptosis) events as well as for the frequency and mean fluorescence intensity (MFI) of Annexin-V^+^ inside the live (PI^-^)/CD11b^+^/ Ly6G^+^ gated populations (see **Supplementary Figure 7** for gating strategy).

### Neutrophil function analysis

Flow cytometric analysis of peroxide production was performed on whole blood leukocytes by quantification of DHR (ThermoFisher, D632) fluorescence upon stimulation with PMA (30 nM) (Sigma-Aldrich, P8139) (additional details in **Supplementary Methods.** See **Supplementary Figure 7** for gating strategy).(Jirapongsananuruk et al. 2003) Flow cytometric measurements of phagocytosis and neutrophil extracellular trap (NET) formation (NETosis) were evaluated on negatively magnetically enriched neutrophils. Flow cytometric quantification of phagocytosis of fluorescently labeled *E. coli* (ThermoFisher, P35366), *S. aureus* (ThermoFisher, P35367) and zymosan (ThermoFisher, P35365) particles followed the manufacturer’s methods for pHrodo™ Green BioParticles® Conjugates (ThermoFisher, MAN0007045) (additional details in **Supplementary Methods.** See **Supplementary Figure 7** for gating strategy).

Flow cytometric quantification of NETosis was performed as previously described.(Gavillet et al. 2015) Briefly, peripheral blood neutrophils, isolated by negative magnetic enrichment, were stimulated with PMA (100 nM) for 4 hours at 37°C, then incubated sequentially with the primary anti-citrullinated histone H3 antibody (Abcam, ab5103), Alexa Fluor700-conjugated secondary antibody (Thermo Scientific, A-21038), and FITC-conjugated anti-myeloperoxidase antibody (Abcam, ab90812), then with Hoechst 33342 (Thermo Scientific, H3570) and analyzed by flow-cytometry (additional details in **Supplementary Methods.** See **Supplementary Figure 7** for gating strategy).

Peripheral blood neutrophil granule content and release were evaluated in negative-magnetically-enriched neutrophils in RPMI 1640 medium with 10% heat-inactivated fetal bovine serum were incubated with PMA (30 nM) for 1 hour to stimulate degranulation. Cell suspension and cell pellet lysates were analyzed for neutrophil elastase (R&D Systems, MELA20) and arginase-1 (Abcam, ab269541) contents by ELISA assays using the manufacturer’s methods.

### Cell Migration

Mice were injected intraperitoneally with 3 mL of 4 % aged thioglycolate solution to induce acute peritonitis; DPBS was injected as a control.(Fukui et al. 2022) Three hours later, animals were euthanized and the peritoneal cavity washed with sterile DPBS. The collected volume was measured and viable cells counted; cell type distributions were determined by automated hematology analyzer (VetScan HM5, Abaxis) and confirmed by flow cytometry.

### Statistics

Descriptive statistics and power analysis sample size evaluations were performed, and the results represented by plots showing mean, interquartile range, overall range and data points. Unpaired Student’s t-test was applied to compare the means of two independent groups. ANOVA and Dunnett’s multiple comparison test were applied to determine which group means were significantly different from their specific control groups, and ANOVA and Tukey’s multiple comparison test were applied to determine which specific pairs of group means differed significantly among multiple groups. A p-value of <0.05 was considered significant. Data analysis was conducted using GraphPad Prism version 10.4.0 (GraphPad Software, Inc.).

## Author contributions

J.S.B., K.N., M.M and P.N. conceived and planned the experiments and interpreted results. J.S.B., Z.Z., K.N., N.B., and M.F. performed experiments and data analysis. P.O-N. contributed to the affinity purification of the anti-Vps45 antibody and optimizations of the immunoblots. B.W. performed the histologic analysis. C.K. developed and studied the *Vps45* knockout mice and VPS45 knockout HeLa cells. J.S.B. and P.N. wrote the manuscript. All authors provided critical feedback and reviewed and approved the manuscript.

## Acknowledgments

We thank the UMass Chan Medical School Transgenic Animal Modeling Core and Dr. Michael Brodsky for their contributions to the generation of *Vps45*^T224N^ and *Vps45*^E238K^ mice; Jennifer Forcina for her contribution to the generation of the anti-VPS45 antibody and preliminary observations of cellular phenotypes; Milka Koupenova, Olga Vitseva and Heather Corkrey for their contributions to the studies on platelets; and Kate Gordon for her contributions to the body health condition scoring analysis.

## Competing interests

The authors have no competing interests to disclose.

## Funding

This work was supported by U.S. National Institutes of Health R01AI152446 (MM, PEN); U.S. Department of Defense BM160046 (MM, PEN) and U.S. National Institutes of Health Training Grant T32-AI095213 (KN).

## Data availability

The data that support the findings of this study are available from the corresponding author upon reasonable request.

## List of abbreviations

Acta2: actin alpha 2
CBC: complete blood count
citH3: citrullinated histone H3
Col3A1: pro-alpha1 chains of type III collagen
DHR: dihydrorhodamine 123
DPBS: Dulbecco’s phosphate buffered saline
G-CSF: granulocyte colony-stimulating factor
HRP: horseradish peroxidase
IMPC: International Mouse Phenotyping Consortium
KI: knock-in
KI/KI: homozygous knock-in mutant
KO: knockout
LPS: lipopolysaccharide
MFI: mean fluorescence intensity
MPO: myeloperoxidase
NET: neutrophil extracellular trap
NETosis: NET formation
PCNA: proliferating cell nuclear antigen
PI: propidium iodide
PMA: phorbol-1-myristate-13-acetate
PCR: polymerase chain reaction
RPMI: Roswell Park Memorial Institute medium
RT-qPCR: real time quantitative PCR
SCN5: severe congenital neutropenia type
5 SCN: severe congenital neutropenia (ss)
DNA: single-stranded donor DNA
VPS45/Vps45: vacuolar protein sorting 45 homolog
*Vps45*^KO^: Vps45 knockout mutant
*Vps45*^T224N^: Vps45 T224N knock-in mutant
*Vps45*^E238K^: Vps45 E238K knock-in mutant
WT: wild-type

## Notes

### Competing Interest Statement

The authors have declared no competing interest.

### Summary of Updates

Corrected spelling of Peter Opara-Nadi's name in the metadata

